# Active Participation of Membrane Lipids in Inhibition of Multidrug Transporter P-Glycoprotein

**DOI:** 10.1101/2020.11.15.383794

**Authors:** Karan Kapoor, Shashank Pant, Emad Tajkhorshid

## Abstract

P-glycoprotein (Pgp) is a major efflux pump in humans, overexpressed in a variety of cancers and associated with the development of multi-drug resistance. Allosteric modulation by various ligands (e.g., transport substrates, inhibitors, and ATP) has been biochemically shown to directly influence structural dynamics, and thereby, the function of Pgp. However, the molecular details of such effects, particularly with respect to the role and involvement of the surrounding lipids, are not well established. Here, we employ all-atom molecular dynamics (MD) simulations to study the conformational landscape of Pgp in the presence of a high-affinity, third-generation inhibitor, tariquidar, in comparison to the nucleotide-free (APO) and the ATP-bound states, in order to characterize the mechanical effects of the inhibitor that might be of relevance to its blocking mechanism of Pgp. Simulations in a multi-component lipid bilayer show a dynamic equilibrium between open(er) and more closed inward-facing (IF) conformations in the APO state, with binding of ATP shifting the equilibrium towards conformations more prone to ATP hydrolysis and subsequent events in the transport cycle. In the presence of the inhibitor bound to the drug-binding pocket within the transmembrane domain (TMD), Pgp samples more open IF conformations, and the nucleotide binding domains (NBDs) become highly dynamic. Interestingly, and reproduced in multiple independent simulations, the inhibitor is observed to facilitate recruitment of lipid molecules into the Pgp lumen through the two proposed drug-entry portals, where the lipid head groups from the cytoplasmic leaflet penetrate into and, in some cases, translocate inside the TMD, while the lipid tails remain extended into the bulk lipid environment. These “wedge” lipids likely enhance the inhibitor-induced conformational restriction of the TMD leading to the differential modulation of coupling pathways observed with the NBDs downstream. We suggest a novel inhibitory mechanism for tariquidar, and potentially for related third-generation Pgp inhibitors, where lipids are seen to enhance the inhibitory role in the catalytic cycle of membrane transporters.

## Introduction

ATP-binding cassette (ABC) transporters form one of the largest transporter superfamilies expressed in cellular and intracellular membranes.^1–3^ They use ATP as a source of energy to actively transport various substrates between the two sides of the membrane. A group of ABC transporters, known as ABC exporters, provide a major mechanism for active efflux of small molecules out of the cell.^4,5^ ABCB1, or P-glycoprotein (Pgp), a prominent member of this superfamily, is ubiquitously expressed in different tissues^6^ where it exports a wide range of charged and neutral hydrophobic molecules.^7^ This promiscuous quality of Pgp is exploited by, e.g., cancer cells that over-express the transporter in their cellular membranes, and thereby acquire resistance to chemotherapeutic reagents and other drug molecules.^8^ In this context, a major drug development effort has been aiming at inhibiting Pgp and therefore achieving more effective chemotherapeutic treatments from the currently used anticancer drugs. In order to make such efforts more effective, it is imperative to understand the mechanism of the transporter in its physiological context, i.e., the cellular membrane. Such understanding will also provide deeper insights into the fundamental questions about the function of ABC transporter and how it may be modulated by the main constituents of the cellular membrane, i.e., lipids, which are present at the immediate vicinity of the protein in a very dense arrangement and can interact with the protein in many different ways.

Structurally, Pgp shows a pseudo two-fold symmetry, folding into a single-chain heterodimeric structure containing two transmembrane domains (TMDs), each consisting of a six-helical transmembrane leaflet associated with a nucleotide-binding domain (NBD).^9–12^ Somewhat common in structurally known ABC exporters, two prominent entry portals exist within the transmembrane region of Pgp, one on either side of the TMDs, which in principle can allow access not only to small molecules to diffuse from within the membrane into the large, central, semi-hydrophobic lumen of the transporter but also to lipids that are frequently observed to bind to these portal regions in the resolved structures of Pgp^10^ and other ABC transporters.^13^

Substrate translocation in ABC transporters like Pgp involves complex structural transitions between inward-facing (IF) and outward-facing (OF) states of the transporter, alternatively exposing the substrate binding site/region to the inside and outside of the cell, a model coined as the “alternate access” mechanims.^14–16^ These conformational changes in ABC transporters are closely coupled to and fueled by binding and hydrolysis of ATP in the NBDs. In Pgp and other ABC exporters, this mechanism facilitates the export of substrate molecules out of the cell.^16–23^ The conformational coupling between the NBDs and the TMDs is key to the ATP-driven transport in Pgp and other ABC transporters. Without getting into specifics, in this transport scheme, ATP binding to the two NBDs promotes their dimerization, with the resulting conformational changes directly coupled to the TMDs and their subsequent transition into the OF state.^18,19^ Hydrolysis of ATP, on the other hand, releases the dimerized NBDs, allowing the transporter to reset to the IF state.^24–26^ Through these general couplings, ATP binding and hydrolysis in the NBDs mechanically control the inter-conversion between the IF and OF, and therefore, the accessibility of the substrate between the two sides of the membrane.

As a multi-domain protein, Pgp exhibits a complex behavior, with the binding of ATP, substrates and inhibitors directly modulating the conformational states and characteristics of the transporter.^27^ Pgp crystal structures, which have been resolved in a non-membrane environment, have mostly found the transporter in excessively open, IF(-like) states, characterized by large separations between the two TMD leaflets on the cytoplasmic side, and between the two NBDs.^9–12^ Similarly, other experimental biophysical studies conducted in non-native environments such as micelles have reached comparable conclusions regarding the open IF state and its highly dynamic nature.^28–31^ On the other hand, biochemical studies performed in more realistic and physiological environments (e.g., nanodiscs or other lipid bilayers), have presented an alternate view of the transporter’s mode of action, where the APO (nucleotide-free) and ATP-bound transporter states are present in a thermodynamic equilibrium capable of sampling similar conformations, ^32–34^ highlighting the importance of the lipid bilayer in regulating Pgp’s conformational dynamics.

A number of studies have also aimed at developing novel inhibitors for targeting this biomedically important transporter.^35^ Out of these, third-generation inhibitors, displaying higher binding affinity and specificity for Pgp, provide the most promising candidates. Tariquidar (TAR), a large (*∼*650 Da), positively charged, amphiphilic compound, is one such high-affinity third-generation inhibitor that binds to the central cavity of Pgp, and it has been suggested to function by destabilizing NBD conformations necessary for ATP hydrolysis.^31^ Recent structural studies have provided additional static information on the location and binding mode of other third-generation inhibitors,^36^ but the molecular details of the mechanism of action of these molecules remain unclear.

Here, we present a molecular dynamics (MD) study providing an atomic-level description of the conformational dynamics of Pgp in the APO, ATP-bound, and TAR-bound states in a natural environment of a lipid bilayer, and propose a model connecting these changes to the possible inhibition mechanisms of Pgp. Our simulations capture a dynamic equilibrium between the open and closed IF conformations in APO Pgp, with ATP binding observed to promote NBD dimerization and closing of the cytoplasmic gate. Compared to the APO and ATP-bound states, TAR-bound Pgp shows highly dynamic NBDs and TMD portal regions, sampling more widely open IF conformations. We also observe a substantially enhanced inhibitor-mediated recruitment of the membrane lipids into the central lumen of the transporter, which together with the inhibitor binding itself result in a differential allosteric coupling between TMD and NBDs, highlighting an important role of lipids in modulating the conformational landscape of Pgp and its inhibition. While the study here focuses on a Pgp inhibitor, the findings regarding the role of lipids can be expected to exist for other types of ligands binding Pgp including the transported substrates, though the mode and degree of lipid interaction with the ligand and the protein can be different in each case.

## Methods

In order to characterize the conformational dynamics of Pgp in the presence and absence of the third-generation inhibitor TAR, we modeled and performed atomistic simulations of 3 different systems, namely, APO (nucleotide-free, drug-free), ATP (ATP/Mg^2+^-bound), and TAR (ATP/Mg^2+^-bound, TAR-bound). We discuss the system construction and the MD methodology in the following sections.

### System Preparation

A recent crystal structure of Pgp (PDB: 4M1M)^12^ in the IF state was used as the starting point for constructing APO, ATP, and TAR systems. For the generation of the APO system, only chain B was retained from the PDB structure, containing residues 34-626 and 696-1,271. The missing region (residues 627-695) corresponds to a flexible linker region connecting NBD1 to the TMD2, not resolved in any available structure of Pgp and was not included in the simulations. For the generation of the ATP system, ATP and Mg^2+^ were docked into their respective binding sites in the two NBDs using a rigorous protocol described by Wen et al,^37^ which reproduces the conserved nucleotide binding characteristics observed in higher-resolution structures of ABC transporters (for this study, the structure of HlyB^38^ was used as the template).

For the generation of the TAR system, we used a variant of ensemble docking, which we term as extended-ensemble docking,^39^ that first samples the conformations of the transporter along its global transition pathway and then utilizes a collection of these conformations representing the entire transition pathway for docking. Briefly, the transition pathway between the ATP/Mg^2+^-bound IF state (modeled above) and the OF structure of Pgp (described in Verhalen et al^23^) was generated using steered molecular dynamics (SMD) along specific collective variables optimized for successfully describing the alternate-access mechanism in ABC-transporters.^40,41^ We note that in doing so, we make the assumption that ATP/Mg^2+^ binding to the IF state is sufficient for the protein to transition to the OF state. The SMD simulation resulted in intermediate structures and conformations along the transition pathway that were then used for our extended-ensemble docking approach. Subsequently, docking of TAR, allowing flexibility around all rotatable bonds, was carried out in the extended-ensemble of the protein containing 50 structures generated along the transition pathway. Finally, the docked structure showing the strongest predicted binding affinity (highest docking score) was selected as the starting structure for the TAR systems. This TAR-bound structure shows a C*α* root mean squared deviation (RMSD) of 4.95 Å with respect to the starting structure of the ATP system. Of importance to our comparative study here, the initial TAR-bound structure is also in an IF state, similar to the APO and ATP initial systems simulated. It should also be noted that a recent DEER study showed that inhibitor might have different effects on the conformational dynamics of the transporter depending on the presence or absence of ATP/Mg^2+^ in the NBDs^31^ (with the former case studied in the present study).

### Membrane Embedding

To study the conformational dynamics of Pgp in APO, ATP/Mg^2+^, and inhibitor-bound states, we carried out multiple all-atom MD simulations for the three systems in explicit membranes including anionic lipids, mimicking conditions used in recent FRET and DEER studies on Pgp reconstituted in nanodiscs.^31,33^ Five independent lipid bilayers containing anionic phospholipids 1-palmitoyl-2-oleoyl-sn-glycero-3-phosphocholine (POPC 40%), 1-palmitoyl-2-oleoyl-sn-glycero-3-phosphoethanolamine (POPE 43%), 1-hexadecanoyl-2-(9Z-hexadecenoyl)-glycero-3-phospho-(1’-sn-glycerol) (PYPG 14%), and 1-palmitoyl-2-myristoyl-1,3-bis(sn-3’-phosphatidyl)-sn-glycerol (PMCL 3%) were constructed using the Membrane Builder module of CHARMM GUI.^42^ Subsequently, each state of Pgp was inserted into the membrane independently and the overlapping lipids removed from the bilayer (Fig. 1).

**Figure 1.**
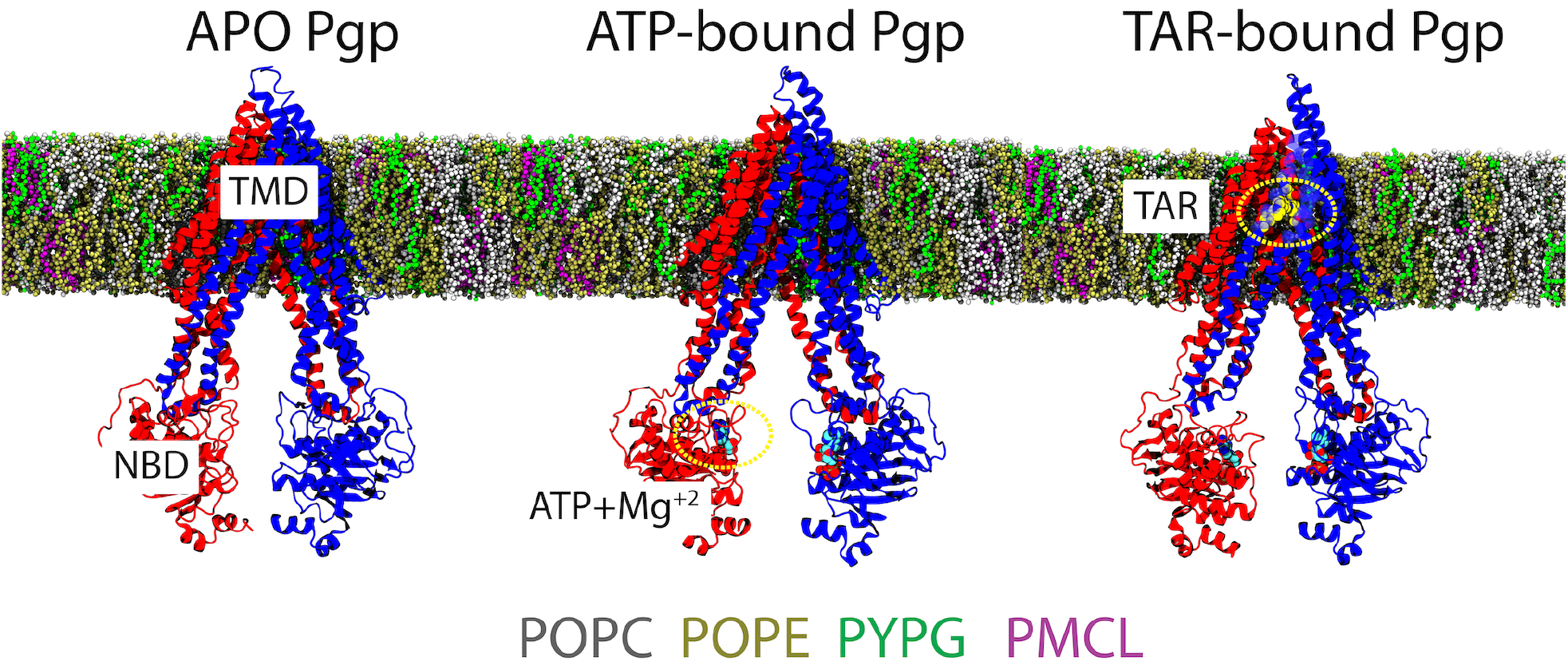
Simulation systems: APO, ATP/Mg^2+^-bound (ATP), and TAR-bound (TAR) IF conformations of Pgp used in the simulations, with the two TMD leaflets, each connected to an NBD, colored in red and blue, respectively (see Methods for details on system construction). All the bound states were inserted in lipid bilayers containing: phosphocholine (POPC=40%), phosphoethanolamine (POPE=43%), phosphoglycerol (PYPG=14%), and cardiolipin (PMCL=3%) lipids (matching experimental conditions in recent FRET and DEER studies on Pgp reconstituted in nanodiscs,^31,33^ shown in different colors. See text for chemical details of the tails.

It is to be noted that in order to remove any bias generated due to the initial distribution of lipids, all the lipids molecules occupying the central cavity of Pgp and/or penetrating the two TMD portals were specifically removed. Thus, two strategies were employed for sampling the conformational space of the studied systems: (A) randomizing the initial distribution of lipids around Pgp, and (B) randomizing the initial velocities of the atoms (Table 1).

**Table 1:** Details of the systems and the simulations performed.

| System name | Ligands bound | No. of independent membrane patches | No. of replicas per patch | Time (ns) /replica |
| --- | --- | --- | --- | --- |
| APO | No ligands | 5 | 2 | 500 |
| ATP | ATP/Mg<sup>2+</sup> | 5 | 2 | 500 |
| TAR | TAR/ATP/Mg<sup>2+</sup> | 5 | 2 | 500 |
| control* | ATP/Mg<sup>2+</sup> | 1 | 1 | 200 |

\* MD simulations performed on TAR-bound Pgp conformation after removing TAR.

In addition to the above-described three systems, we also performed simulations of the starting TAR-bound Pgp conformation after removing the inhibitor. This was done so as to check the stability of the starting TAR-bound Pgp conformation, as well as a point of comparison to the starting conformations of the APO and ATP-bound systems. We term this structure as the “TAR-less” system or the “control” simulation.

### MD simulations

All the simulations were performed using `NAMD2.`^43,44^ CHARMM36m protein and lipid force field^45,46^ and TIP3P water^47^ were used. The inhibitor molecule (TAR) was parametrized using the `force field ToolKit (ffTK)` plugin^48^ of `VMD`.^49^ Briefly, the geometry optimization was performed using second order Moller-Plesset (MP2) theory^50^ and the 6-31G(d) basis set in Gaussian09.^51^ After geometry optimization, partial charges were assigned using quantum mechanical calculations. The bonds and angular terms were calculated directly from the Hessian, and the dihedral parameters were fit to the QM potential energy landscape.

The protonation states of the ionizable residues of Pgp were calculated using the `PropKa` server.^52^ The C-terminal carboxylate capping group, N-terminal ammonium capping group, and all the hydrogen atoms were added to the structures using the `PSFGEN` plugin of `VMD`.^49^ All systems were solvated and neutralized with 0.15 M NaCl. The final systems contained *∼*250,000 atoms each with approximate dimensions of 135*×*135*×*190 Å^3^.

All the simulations were performed as NPT ensembles, at a temperature of 310 K maintained using Langevin thermostat with a damping coefficient of *γ* = 1 ps^−1^. Constant pressure was maintained at 1 bar using the Nosé-Hoover Langevin piston method.^53^ The particle mesh Ewald (PME) method^54^ was used in calculating the long-range electrostatic forces. A 12-Å cutoff distance and a 10-Å switching distance were used in calculating the non-bonded interactions. All the simulations were performed with the keyword `vdwForceSwitching`, in order to use force-based cutoffs. An integration timestep of 2 fs was used in all simulations. The system was initially energy-minimized and equilibrated for 10 ns with the C*α* atoms restrained to their initial positions with a force constant *k* = 1kcal/mol/Å^2^. All restraints were then released in the following 500-ns production run. Additionally, a 200-ns control simulation of the TAR-less system was also carried out (Table 1).

The initial and final structures for all the simulations, topology/parameter files, and configuration files for the runs are all made available in Open Science Framework, https://osf.io/3kyx2/?view_only=54eccc30d54347b0b857d18f7d402df9. Trajectory files will be made available upon request.

### Analysis

The conformational dynamics of Pgp were characterized in terms of the large-scale motions of the different domains of the transporter, as well as secondary structure changes in the different simulated systems. As the surrounding lipid environment can play a significant role in modulating the conformational ensemble of membrane proteins, we characterized the lipid density and penetration inside the protein in the different systems, as well. Additionally, we also characterized the potential conformational coupling between the TMD and NBDs of Pgp, as the structural changes observed in the different systems may differentially affect the communication between these domains. Analysis on the conformational changes in the protein and calculation of lipid properties around the protein were carried out on all the simulation runs. All analyses were performed using our in-house scripts and/or VMD plugins.

#### Global conformational dynamics

ABC transporters, and in particular ABC exporters, are highly dynamic systems undergoing complex structural changes in both their TMDs and NBDs. The global conformational dynamics of Pgp in all the simulated systems were monitored based on a set of orientational collective variables that have been previously used to successfully describe the alternate-access transport model^14^ in homologous ABC transporters.^40,41^ Briefly, the angle *α* is defined by the relative orientation of the two TMD leaflets and describes the cytoplasmic opening of the TMDs. The angle *γ* is defined by the relative orientation of the two NBDs and describes their relative twist as they approach each other (Fig. 2A). Additionally, distances were calculated between different domains of the transporter: (a) between the C-*α* atoms of residues S92 and R745 located at the extracellular side of Pgp (d_*ext*_) to represent the opening of the extracellular mouth of the transporter, (b) between the C-*α* atoms of residues K145 and M787 located in opposite leaflets (d_*cyt*_) to represent the separation of the TMDs on the cytoplasmic side, and (c) between the centers of masses of the two NBDs describing their degree of dimerization (d_*NBD*_) (Fig. 2A). These orientational and distance-based metrics together capture major conformational changes in the TMDs and the NBDs of Pgp.

**Figure 2.**
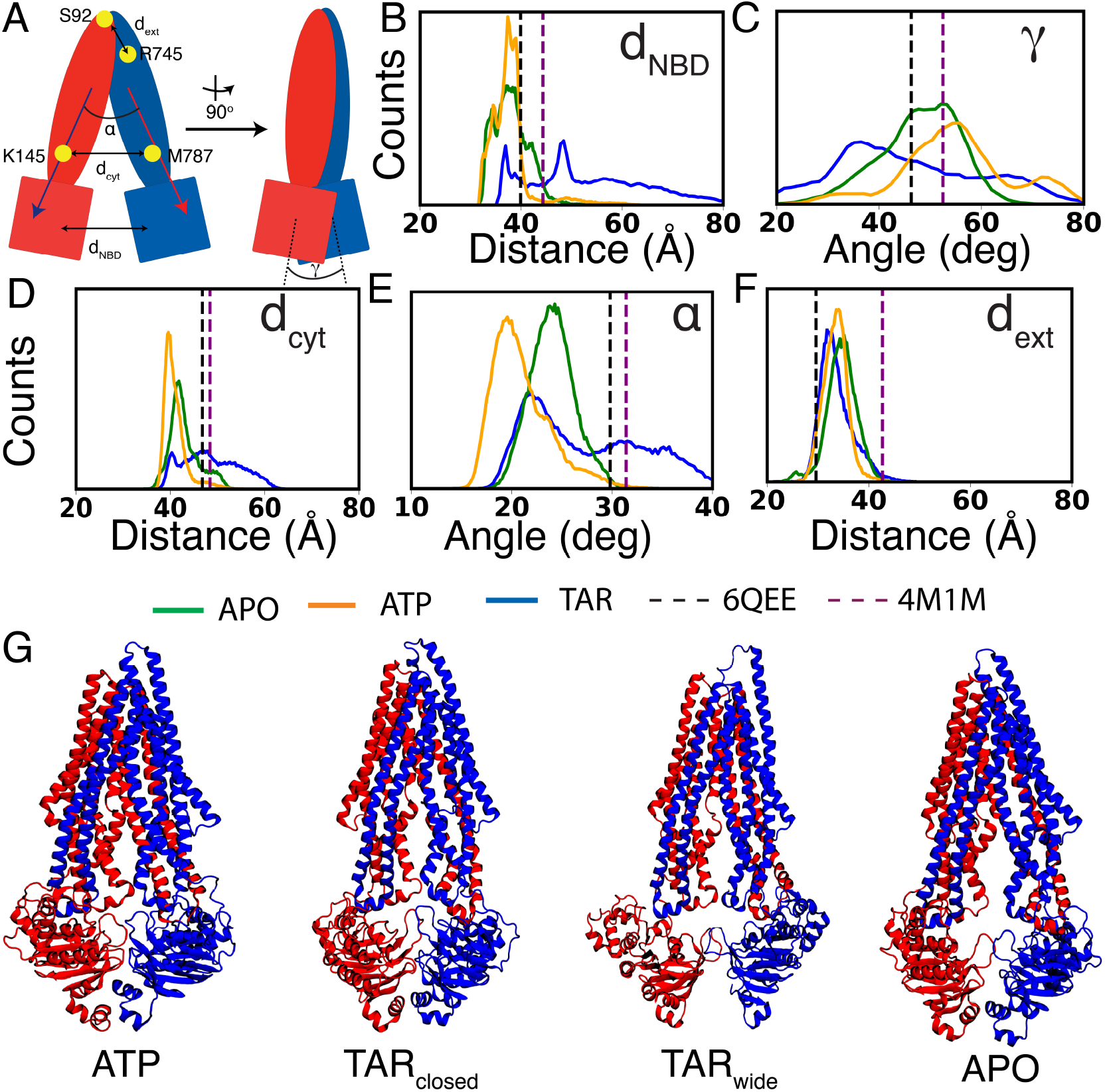
Conformational dynamics of Pgp in different bound states: (A) Schematic representation of Pgp depicting the geometric parameters used for monitoring its global conformational dynamics. The two halves of the protein are colored in red and blue. Conformational dynamics in all the simulated systems were evaluated by analyzing: (B) d_*NBD*_: the center of mass distance between the two NBDs, (C) *γ*: relative twist of the two NBDs, (D) d_*cyt*_: the distance between C*α* atoms of K145 and M787 on the cytoplasmic side of the TMDs, *α*: the angle describing relative orientation of the TMD helices measuring the degree of cytoplasmic opening, and, (F) d_*ext*_: the distance between C*α* atoms of S92 and R745 on the extracellular end of the TMDs. The histograms in B-F are calculated using the entire dataset and indicate the presence of more widely open IF conformations in the TAR-bound system (shown in blue), compared to APO (green) and ATP (orange) systems. Dotted vertical lines in each panel mark the values in the recent structures for an inhibitor (zosuquidar)-bound Pgp (PDB:6QEE)^36^ (black) and for APO Pgp (PDB:4M1M)^12^ (purple). Time series of the distances shown in the histograms are provided in Fig. S1. (G) Representative snapshots for ATP, TAR_*closed*_, TAR_*wide*_, and APO are shown.

#### Local conformational dynamics

The recently reported structures of Pgp display differences in their secondary structure, especially in their TMDs. ^36,55^ In addition to the global conformational changes, we thus quantified the local changes (defined for the immediate region surrounding the central cavity where the inhibitor binds) in the different simulated systems. In doing so, we calculated the root mean square fluctuations (RMSF) of individual residues as well as the secondary structure content of the portal helices (TM4, TM6, TM10, and TM12). Both these calculations were performed using VMD.

#### Lipid density around the protein

The surrounding lipid environment can have a significant effect on the structural dynamics of membrane proteins. The role of the surrounding lipidic environment was thus evaluated by calculating the lipid distribution around Pgp in all the simulated systems. The lipid density profiles were calculated by plotting the 2D histogram of the position of the phosphorus atoms in all the simulated replicas for each system.

#### Lipid access to the lumen of Pgp through the TMD portals

Differences observed in lipid density around the protein in different conformational/bound forms, as discussed later, also pointed to possible differences in lipid access inside the protein’s lumen. We therefore calculated the number of times lipids penetrated into the the central cavity of Pgp (lipid access events) for all the three studied systems, as described below. The central binding region of Pgp is formed by residues from the TM1, TM2, TM3, TM5, TM7, TM8, TM9, TM11, and TM12 helices. A lipid access event was counted if the phosphate head group of a lipid molecule penetrated entirely the cylindrical region defining this cavity at any point during the simulation. To characterize the movement of the lipids that access the Pgp central binding region/cavity, the *z* components of the centers of masses of individual lipid head groups with respect to the membrane midplane (lying at *z* = 0) were monitored over the course of the simulations.

#### >Lipid-interacting residues in TMD portals

The lipid-protein interaction analysis was performed to identify the portal residues forming the hotspots for lipid interactions on the surface of Pgp. The entry portals are delimited by the helices TM4 and TM6 in Portal 1, and by TM10 and TM12 in Portal 2. An interaction between a lipid and a portal was defined whenever the phosphate head group of the lipid molecule came within 3 Å of a portal residue. The heavy atoms of the following portal residues were considered for this analysis: K230-H241 of TM4, S345-G356 of TM6, L875-K883 of TM10, and Y994-S1002 of TM12. These calculations were done for the entire trajectory, yielding the probability of a residue forming contacts with the lipid molecules.

#### Network analysis of TMD-NBD coupling

The transport cycle of ABC transporters involves large conformational changes in both TMDs and NBDs, which rely on complex communication pathways between the two parts of the protein. Coupling between TMDs and NBDs can somewhat be evaluated in terms of the pathway of residues that facilitate the mechanical coupling, and therefore mediate force propagation between the two domains. To investigate the inter-domain communication, dynamic network analysis was performed on different Pgp systems using the Network-View plugin^56^ of VMD. In a network, all C-*α* atoms are defined as nodes connected by edges if they are within 4.5 Å of each other for at least 75% of the MD trajectory. Pearson correlation was first used to define the communities in the entire network corresponding to sets of residues that move in concert with each other. Furthermore, the coupling between the TMDs and NBDs was defined as the path connecting a source and a sink. The source was defined at the site of residue S988, located on the portal helix TM10 that shows major conformational variability between the three simulated bound states. The sink was defined by Q1114 of the conserved Q-loop of the NBD, making contacts with the coupling helices in the TMDs and important for coupling conformational changes between the NBDs and the TMDs.^57^ The tolerance value used to include a path in the suboptimal pathway was -ln(0.5) = 0.69. The network analysis was performed taking into account the full length of the 500-ns MD trajectories.

#### Structural assessment of the inhibitor’s docking pose

In order to further validate the stability of the starting binding mode of TAR in the Pgp central cavity, as captured from our extended ensemble docking studies, we monitored two complementary properties: (1) the RMSD of TAR with respect to Pgp from the 500-ns simulations, and (2) the contacts between the heavy atoms of TAR and its binding pocket residues. A distance cut-off of 3.5 Å was used to calculate these contacts. Furthermore, we compared these contacts with a recent cryoEM structure of Pgp in complex with another high-affinity, third generation inhibitor, zosuquidar.^58^

## Results

Multiple all-atom MD simulations of Pgp in APO, ATP- and inhibitor-bound states allowed us to characterize major structural and dynamical differences between these states, also highlighting the influential and differential role played by the lipids in the dynamics of the transporter. The results presented here are primarily divided into three sections describing: (1) global and local conformational dynamics of Pgp in different bound states, (2) lipid-mediated structural modulation of Pgp, and, (3) allosteric coupling between the TMDs and NBDs.

### Conformational dynamics of different bound states of Pgp

#### Global conformational dynamics

The global conformational dynamics of Pgp in the APO, ATP and TAR systems were characterized in terms of the relative orientation and the distances of different domains. The TMD conformational dynamics in Pgp were described by the angle *α*, defining the degree of opening on the cytoplasmic side, and inter-residue distances calculated at the base as well as apex regions of the TMDs (d_*cyt*_ and d_*ext*_, respectively). The degree of NBD dimerization was described in terms of the twisting angle between the NBDs, *γ*, and the respective separation between their centers of masses (d_*NBD*_) (Fig. 2A). During the simulations, the ATP systems show significant closing and more frequent dimerized-like states of the NBDs compared to the starting crystal structure (Fig. 2B,C,G and S1). On the other hand, the inhibitor-bound (TAR) systems show highly dynamic NBDs, both in terms of the distance and the twist angle between them, sampling two distinct conformational basins defined based on their NBD separation: TAR_*wide*_ conformations characterized by d_*NBD*_ values larger than 42 Å and TAR_*closed*_ conformations if the distance is shorter (Fig. 2). In the APO systems, we observe a dynamic equilibrium between the open- and closed-IF conformations, and relatively higher populations for the open NBD conformations compared to the ATP systems.

Similar to NBD distances, a significant degree of cytoplasmic TMD closing (d_*cyt*_) is observed for the ATP systems, whereas the APO systems show relatively more open cytoplasmic TMD conformations (Fig. 2A,D and S1). The TAR systems sample a broader range of d_*cyt*_, lying on either side of the starting crystal structure. In terms of the cytoplasmic opening, (*α*), the ATP systems show smaller TMD angles as compared to APO (Fig. 2E). The TAR systems show TMD cytoplasmic angles similar to the APO systems, as well as a large population similar to the starting crystal structure and the recent cryoEM structure of zosuquidar-bound Pgp.^58^ We note that in all the simulated systems, including TAR_*wide*_ and TAR_*closed*_, TAR remains stably bound inside the central cavity of Pgp (Fig. S2). Additionally, comparison of the TAR-bound simulations with the zosuquidar-bound Pgp structure shows a large similarity between the protein residues involved in interactions with the two inhibitor molecules. Pgp may thus display a shared (conserved) binding site for these two molecules, and possibly other third-generation inhibitors in general.

Given that during the simulations, Pgp does not sample conformations in the occluded or OF states, no significant changes are noticeable in the extracellular opening, as measured by d_*ext*_ (Fig. 2F). The control (TAR-less) simulation (the TAR-bound conformation after removing the inhibitor) displays a dynamical behavior similar to the ATP-bound Pgp, supporting the structural similarity of the two protein conformations used to initiate these two simulations (Fig. S3).

#### Local conformational dynamics

The differences in local fluctuations for the three states were characterized by calculating the RMSF of individual protein residues. In inhibitor-bound (TAR) systems, NBD residues were observed to be more dynamic when compared to the APO and ATP systems (Fig. S4A, B, C and D). Interestingly, the RMSF calculations also highlighted that NBD2 is relatively more dynamic compared to NBD1 in the presence of ATP (for both TAR and ATP systems) (Fig. S4A). This asymmetry in the dynamics of the NBDs, not observed in the APO-system, appears to be further amplified in the presence of the inhibitor. Additionally, residues in the coupling helices (156-161, 257-263, 796-800, and 900-905) as well as the portal helices TM4 and TM6 comprising Portal 1, and TM10 and TM12 comprising Portal 2, lying on the opposite sides of the TMD, are significantly more dynamic in the TAR systems compared to the ATP and APO systems. In order to further evaluate the structural dynamics of the portal helices, which show relatively large fluctuations, their secondary structure content was monitored over the course of the simulations. TM10, initially present as a structured helix, was found to be partially unfolded in the inhibitor-bound simulations (Fig. 3A, B). A similar, broken TM10 helix is also captured in the recent cryoEM structure of zosuquidar-bound Pgp.^36,58^ This helix maintains its helicity in ATP simulations, whereas the APO systems show an average helicity between the ATP- and inhibitor-bound systems (Fig. 3A, C and D). Except for TM12, which is partially unfolded in the starting crystal structure, no further unfolding was observed in other portal helices (TM4 and TM6) (Fig. S5).

**Figure 3.**
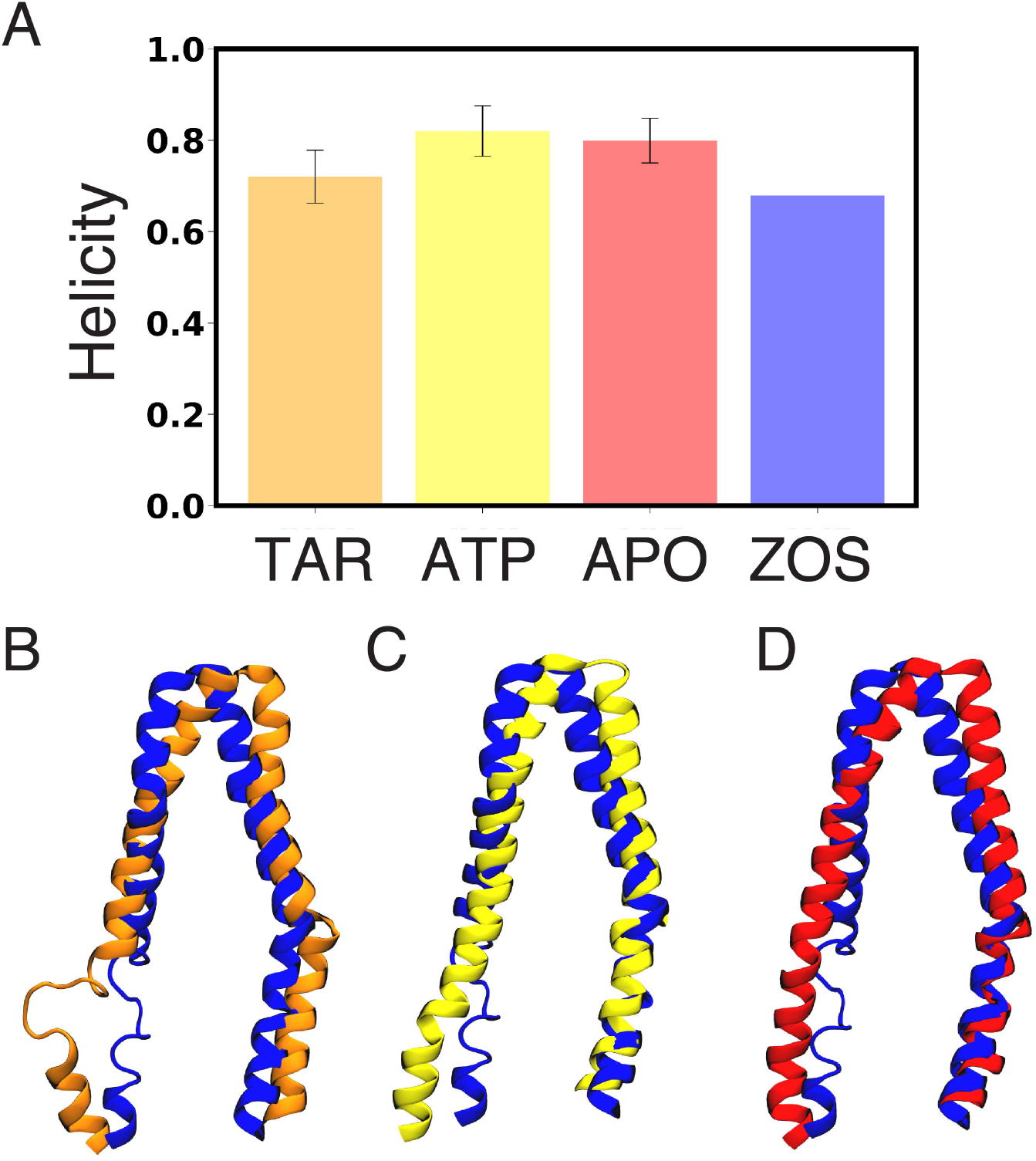
Differences in secondary structure: (A) Comparison of the degree of helicity in the portal helix TM10 (residues 876-889) in APO, ATP, and TAR Pgp with the zosuquidar-bound (ZOS) Pgp cryoEM structure.^58^ The analysis was performed combining all the simulations for each state. The structure of TM10 in a representative trajectory snapshot from: (B) TAR-bound (orange), (C) ATP (yellow), and, (D) APO Pgp (red) aligned with zosuquidar-bound Pgp (ZOS)^58^ (blue). Helicity of other portal helices (TM4, TM6, and TM12) is shown in Fig. S5. A p-value of 0.0 was obtained on comparing the degree of helicity in the different systems using one-way ANOVA test, pointing to statistical significance of the results.

### Lipid-mediated structural modulation of Pgp

The role of membrane lipids in modulating the structural properties of Pgp was further evaluated by calculating the lipid distribution and packing around the protein as well as lipid penetration/translocation inside the central cavity of Pgp.

### Lipid packing around Pgp

To probe the role of the surrounding lipidic environment in modulating the global and local dynamics of Pgp, we first calculated the distribution of lipids around the transporter from our simulations. In the inhibitor-bound systems, a large density of lipids was observed in the central cavity, particularly lipids that originate from the cytoplasmic leaflet and diffuse through the two TMD entry portals (Fig. 4A). The APO systems also show lipids inside the central cavity, though to a smaller degree compared to the TAR systems. In the ATP systems, on the other hand, a considerable lipid density is only observed inside Portal 1 (Fig. 4B and C). All lipids accessing the central cavity in the simulated systems originate from the cytoplasmic leaflet of the membrane, in agreement with the results of a previous coarse-grained simulation study,^59^ and clearly related to the more open configuration of Pgp in the cytoplasmic half.

**Figure 4.**
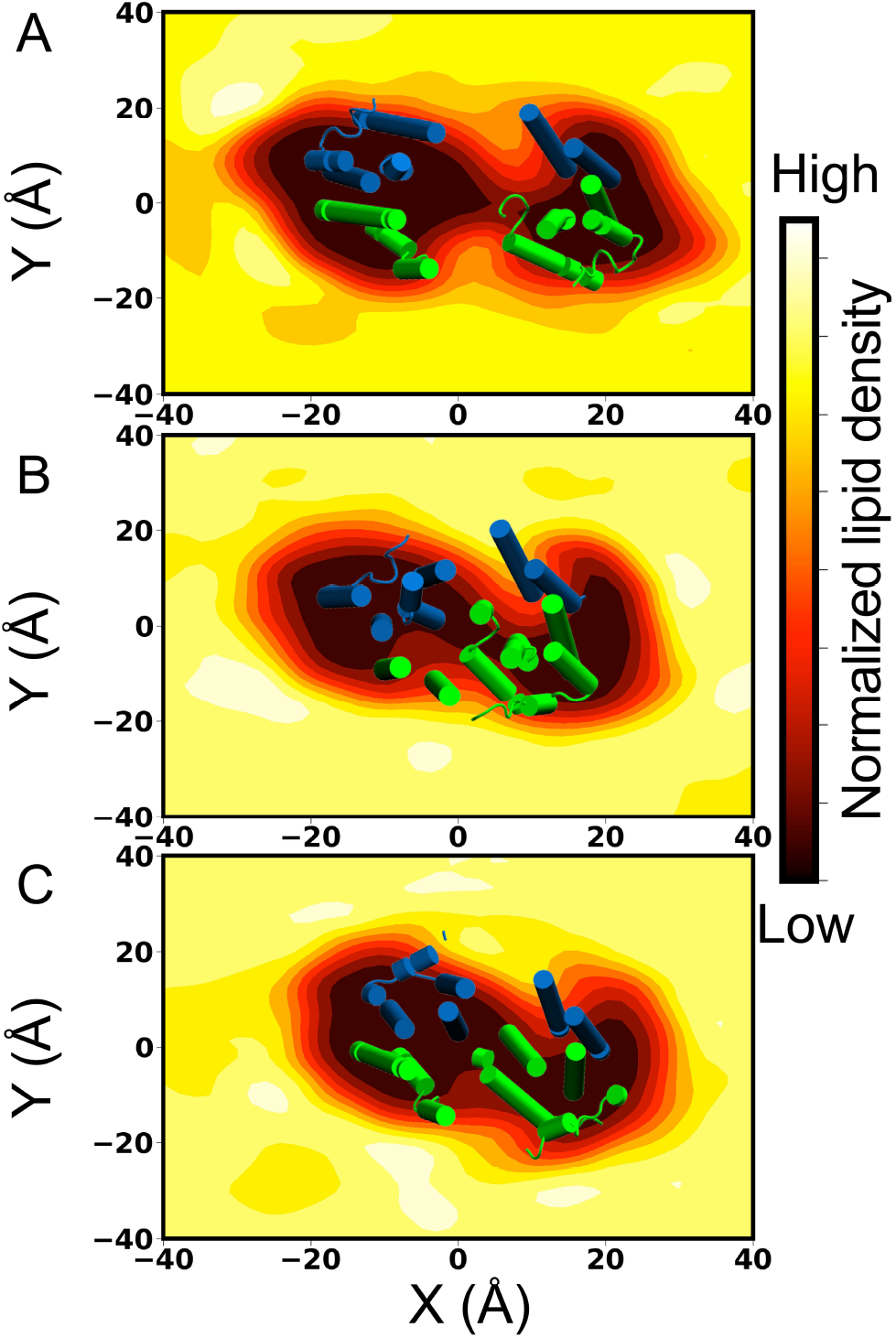
Lipid packing around and inside Pgp: The cytoplasmic-leaflet lipid density profile for (A) TAR, (B) APO, and (C) ATP Pgp simulations was calculated over all the simulated replicas for each system. The molecular systems are aligned along the *Z*-axis and viewed from the extracellular side of the protein, with the *XY* indicating the plane of the membrane. All histograms are normalized the same way, that is, with respect to the average bulk density of lipids. The lipid molecules access the central cavity through the entry portals located on either side of the TMD. TAR-bound and APO systems show enhanced lipid densities inside the central cavity of Pgp, while a low density was observed in the case of ATP-bound systems. In each panel, a part of Pgp spanning the cytoplasmic membrane leaflet (taken at the end of one of the 500-ns simulations for each system) is shown in cartoon representation with the two Pgp halves colored in green and blue, respectively.

### Lipid access through TMD portals

The number of lipid access events in the simulated systems was calculated by monitoring the positions of lipid head groups assessing the central cavity of Pgp, defined as a cylindrical region (Fig. 5A), over the course of the simulations. As expected, the maximum number of lipid access events is observed for the inhibitor-bound systems, especially for TAR_*wide*_ conformations, and the minimum number of events for the ATP systems (Fig. 5B). Consistent with the higher lipid densities observed inside the central cavity for the APO systems compared to the ATP systems, lipid access events were also found to be relatively lower in the latter case.

**Figure 5.**
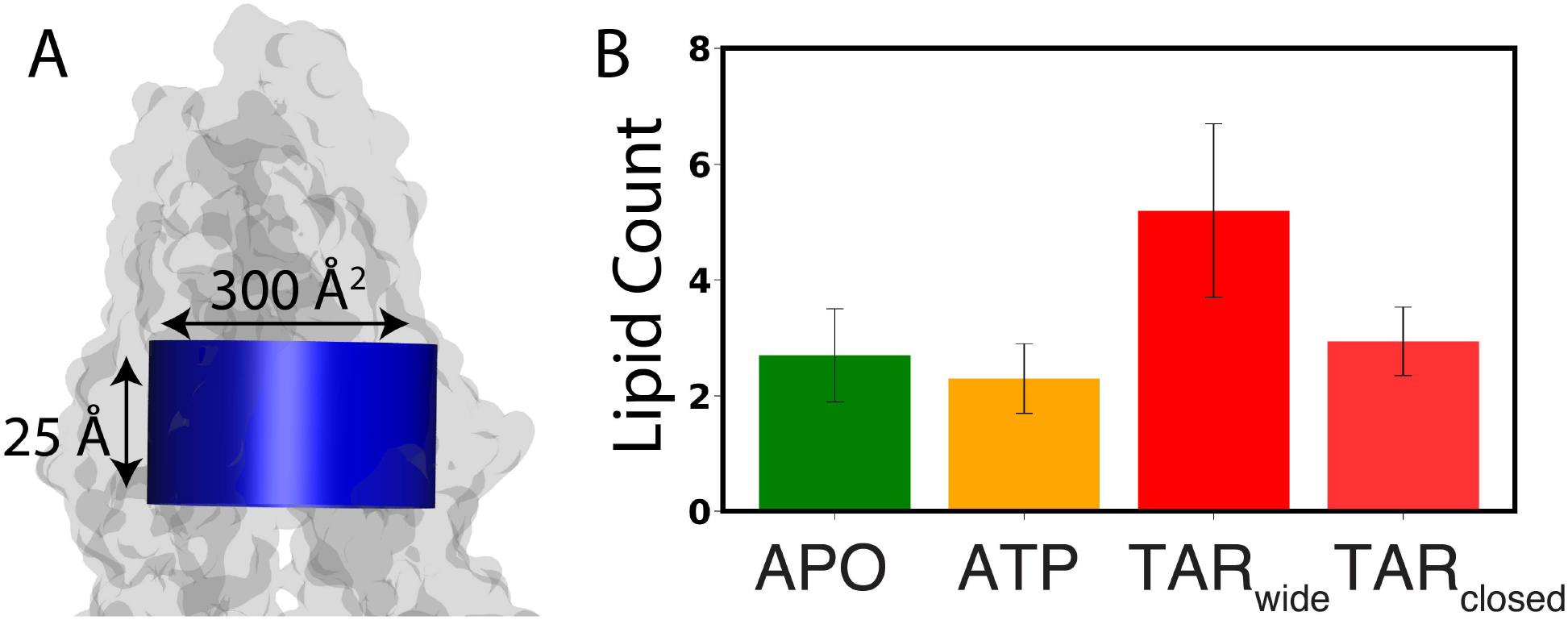
Lipid access into the lumen of Pgp: (A) A lipid access event was counted if the entire head-group phosphate of any lipid accessed the central cavity of Pgp, defined by a cylinder (shown in blue) with a cross-sectional area of 300 Å^2^ and a height of 25 Å (see Methods for the criteria used to define the boundaries of the central cavity). Pgp is shown in a translucent surface representation. (B) Count of lipids accessing the central cavity of Pgp through the entry portals in APO (green), ATP (orange), and TAR (red) simulations, calculated by counting the number of phospholipid head groups accessing the central cavity over the course of the 500-ns simulations. Two TAR-bound conformations (TAR_*wide*_ and TAR_*closed*_) are defined based on the d_*NBD*_ separation (Fig. 2B). In comparison to the APO and ATP systems, more lipid access events are observed in the TAR systems, especially for the TAR_*wide*_ conformations. A p-value of 0.0 was obtained on comparing the lipid access events for the different systems using one-way ANOVA test, pointing to statistical significance of the results.

The key Pgp residues involved in lipid recruitment through Portals 1 and 2 were determined by monitoring the portal residues which make contacts with the lipid head groups (Fig. S6A,B,C). Both entry portals are observed to be rich in basic residues that directly interact with the negatively charged phosphate groups of lipids: R355, K238, and K230 at Portal 1, and K881 and K996 at Portal 2 (Fig. S6D,E,F,G). Interestingly, we also observed system-dependent opening and re-arrangement of the basic residues in Portal 2, which may play a role in the differential lipid recruitment observed in the different systems.

### Translocation of lipid molecules within the lumen of Pgp in TAR-bound systems

To evaluate the behavior of lipids that access the Pgp binding cavity, the membrane positioning (*z* coordinate) of the lipid head groups in the central cavity was monitored over the course of the simulation. Interestingly, in TAR_*wide*_ conformations, significant translocation of lipid head groups along the membrane normal (partial flipping events) were observed. During these events, lipid head groups from the cytoplasmic leaflet translocated along the membrane normal towards the extracellular leaflet (see one full flipping event in Supplementary Video 1), forming direct interactions with the terminal methoxy groups of the bound inhibitor in the central cavity (Fig. 6A and B). The lipids translating beyond the membrane midplane form close and stable interactions with TAR during the remainder of the simulation time. The lipid tails of these ‘wedge-lipids’ remain largely in the bulk membrane connectingto their inserted head groups through the entry portals (Fig. 6B). In total, we captured 12 partial lipid flipping events in the TAR systems, including PE, PC, and CL lipids (Fig. 6A). In contrast to the TAR systems, cytoplasmic lipids that penetrate the Pgp lumen in the APO or ATP systems remain largely within their original leaflet during the simulations (Fig. 6C, D and S7) as well as in the control simulation after removing the bound inhibitor. Given the insufficient number of partial lipid flipping events in our simulations we can not reliably assess any lipid preference.

**Figure 6.**
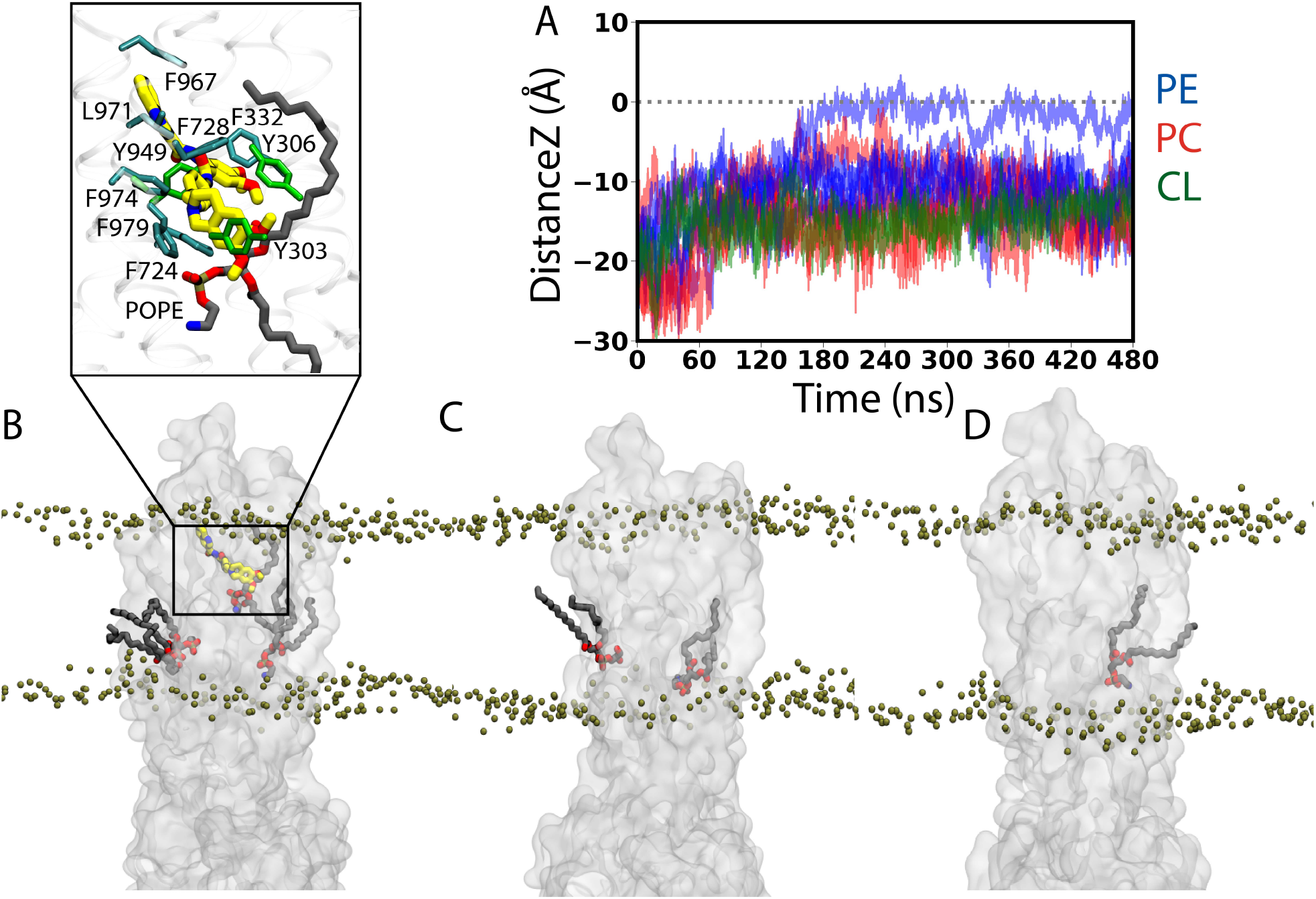
Trans-leaflet lipid translocation inside the lumen of Pgp: (A) Translocation of lipids in the central cavity of Pgp that might result in their flipping was monitored by calculating the *z* component of the center of mass of the lipid head groups in all the simulated systems. In total, we captured 12 lipid molecules from the cytoplasmic (lower) leaflet that undergo partial flipping during the simulations. Data for the APO Pgp are shown in Fig. S7. (B-D) Lipids (licorice representation) penetrating into the central cavity of (B) TAR-bound, (C) APO and (D) ATP Pgp (translucent surface representation) from representative snapshots of the simulations. Phosphorus atoms from the lipid head groups are shown as tan beads to delineate the membrane boundaries. Inset in (B) shows the closeup view of the interactions between the bound TAR (thick sticks) and the flipping lipid (PE) molecule showing the largest translation (black trace in (A)) into the other side of the membrane, as well as the surrounding Pgp residues within 3.5 Å of either TAR or the head group of the translocating (PE) lipid. A 2D representation of the TAR binding pocket along with the flipping PE molecule is shown in Fig S2B.

### Conformational coupling between TMDs and NBDs

Dynamic network analysis was employed to evaluate the allosteric coupling between the NBDs and TMDs in the different simulated systems. The coupling pathway was calculated between TM10 in the TMD, which shows major conformational differences between the three systems (as described above), and the conserved Q-loop in the NBD, which is important for coupling conformational changes between the NBDs and TMDs and subsequent conformational transition between the IF and OF states.^57,60^

The APO and ATP systems predominantly show short and direct coupling pathways connecting the TM10 with the Q-loop (Fig. 7A and B). In the case of the ATP systems, the pathway connects the TMD with the NBD directly through the coupling helices located at the interface of TMD and NBD, whereas in the APO systems, the pathway connects the two domains without directly passing through the coupling helices.

**Figure 7.**
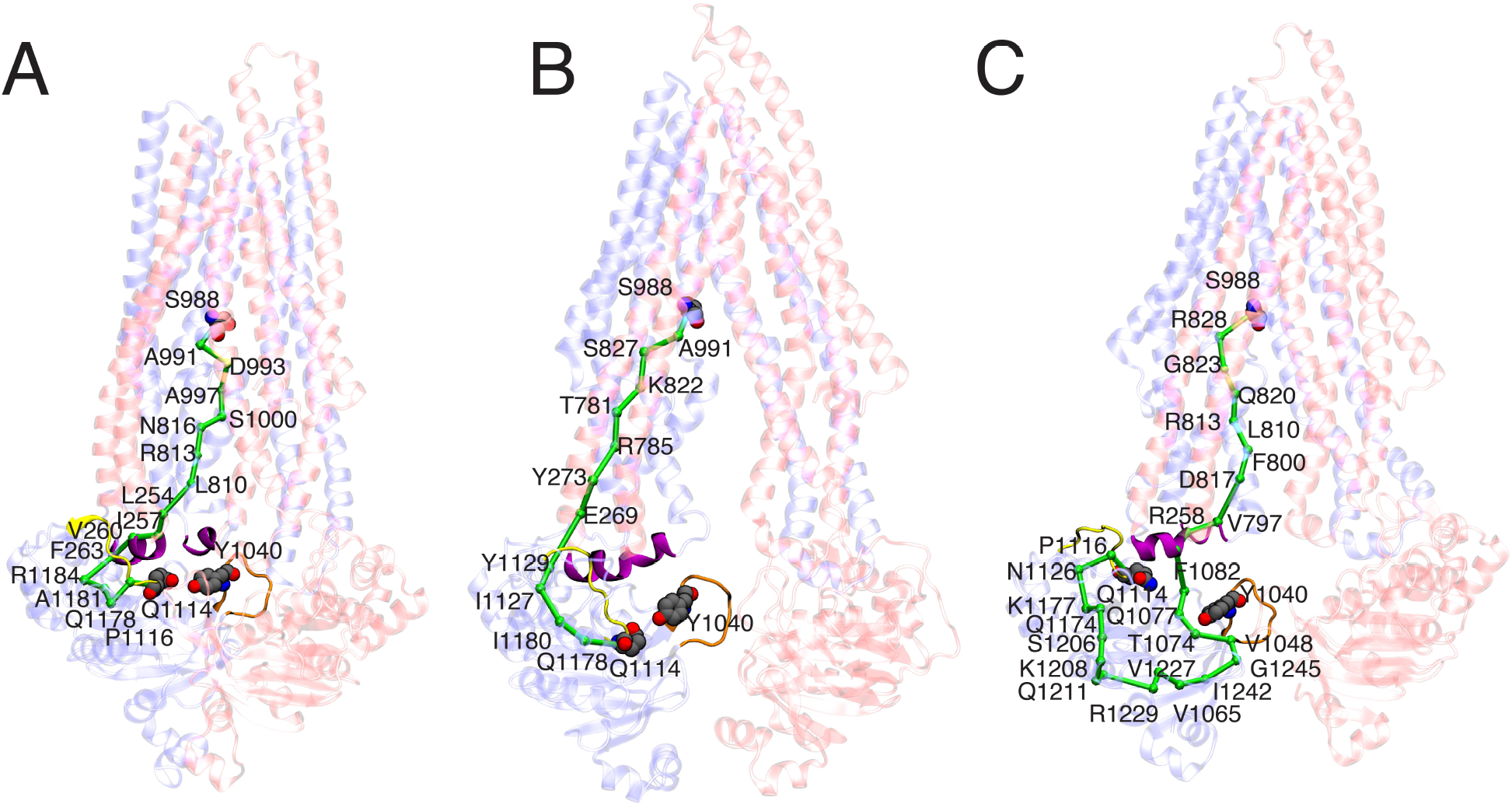
Allosteric coupling between TMDs and NBDs: The coupling pathway calculated for a representative 500-ns long simulation of: (A) ATP, (B) APO, and (C) TAR-bound Pgp, between S988 in TMD (the source) and Q1114 in the NBD (the sink), is shown in green. Q-loop (containing conserved Q1114) is shown in yellow, coupling helices in purple, A-loop (containing conserved Y1040) in orange, and the rest of the protein in a transparent cartoon representation. All the residues that form the allosteric pathway are highlighted in the representative replica for each system. In the majority of TAR-bound simulations, extended coupling pathways passing primarily through the coupling helices and the Q-loop in the NBDs are observed. Shorter and more direct pathways connecting the TMD and the NBD are observed in the ATP and APO simulations.

In the case of the inhibitor-bound systems, on the other hand, elongated pathways are predicted passing through additional portions of the NBD (Fig. 7C). In some cases, the pathway connects the coupling helices with the A-loop, which contains a conserved aromatic residue (Y1040) critical for ATP binding.^61^ As a result, the pathway transverses a larger portion of the NBD before reaching the Q-loop. In addition, by employing dynamic network analysis, we also investigated the communities, i.e., the sets of residues that move in concert with each other during the simulations. Similar communities were observed in the TMDs and the NBDs for all three systems (Fig. S8).

## Discussion

Pgp is a biochemically well studied ABC exporter that actively extrudes certain xenobiotics and other amphiphiles across the plasma membrane. The transporter is overexpressed in a variety of cancer cells where it is closely associated with the development of multidrug resistance by interfering with the therapeutic accumulation of the drugs inside the affected cells. As such, a major effort is invested in developing novel strategies for targeting and inhibiting Pgp. In the present study, we have carried out an extensive set of all-atom MD simulations to characterize the conformational dynamics of Pgp in a number of native states arising during the transport cycle (APO and ATP-bound), as well as in the presence of a third-generation inhibitor, TAR. We capture differential conformational dynamics, both globally and locally, associated with the different bound states of Pgp, and, furthermore, we propose a novel lipid contribution to the mechanism of action of the inhibitor, which might also accompany the binding of other ligand types to Pgp and otehr ABC transporters, though with different modes and in different extents.

The global orientational dynamics captured in our MD simulations of Pgp embedded in multi-component lipid bilayers, each starting from a different lipid distribution around Pgp, show a dynamic equilibrium between open and closed (dimer-like) NBD conformations of the transporter in the APO system, with the NBD twist angles showing large variations around the crystal structure values (Figs. 2 and Fig. S1). Along with NBD dynamics, the transporter also samples distances corresponding to both more open and more closed TMDs in comparison to the starting crystal structure used in the simulation, but it remains relatively open with respect to the ATP-bound state. The APO systems also show a higher lipid density inside the binding cavity of Pgp as well as a greater number of lipid access events than the ATP systems, which both can be attributed to the more open configuration of the TMDs on the cytoplasmic side (Fig. 4 and Fig. 5). As the substrates are believed to access the central binding cavity of Pgp largely from the lipid bilayer,^32^ the larger cytoplasmic TMD opening in the APO state is expected to provide an ideal pathway for this process. In this context, the pathway observed in our simulations for the penetrating lipid molecules accessing the central cavity of Pgp may also represent a potential route for the lateral translocation and diffusion of the substrate molecules into the central cavity. The excessively open, IF-like structures of APO Pgp that have been observed in structural studies using reconstituted Pgp in detergent or micellar environments,^9–12^ confer it with no added physiological function and may portray artifacts of the non-physical (non-membrane) environments used in these experiments. The more closed Pgp conformations captured in the APO simulations, on the other hand, have been recently characterized in cryoEM,^34^ luminescence resonance energy transfer (LRET),^33^ as well as atomic-force microscopy experiments (AFM),^32^ when Pgp was reconstituted in lipid nanodiscs or in liposomes, constituting a more natural membrane environment for the transporter.

In the ATP-bound systems, the conformational equilibrium is shifted towards closed-IF like states characterized by closure of the TMD cytoplasmic gate in addition to dimerization of the NBDs (Fig. 2 and Fig. S1). The ATP systems in our simulations lie close to the recent ATP-bound dimerized structure of human Pgp,^55^ adopting a closed NBD conformation coupled to a closed TMD. As expected, a low lipid density is observed within the central cavity of Pgp in the ATP systems which can be attributed to the partial closure of the cytoplasmic side of the TMDs in these systems (Fig. 4 and Fig. 5). Substrate binding and subsequent binding of ATP to the NBDs in the APO systems may concurrently drive occlusion of the substrate-bound TMD and the complete closure of the inter-NBD gap, ^36^ in line with the conclusions of earlier mutagenesis and biochemical studies.^33,62^ Alternatively, ATP binding prior to the substrate entry may drive the transporter to a substrate-free occluded state, responsible for the known basal ATPase activity of the transporter.^63,64^ A close interaction between the coupling helices in the TMDs and the Q-loop in the NBDs, observed more in the ATP-bound systems than the APO systems (Fig. 7), may allow fine-tuning of the coupled movements of different domains of the transporter necessary for the transition to the OF state and completion of the transport cycle, events which are not fully observed here due to the limited timescale of the simulations.

From our systematic MD study, clear differences are observed in the conformational ensemble of Pgp in the presence of an inhibitor bound to the central cavity. In the presence of the inhibitor, TAR, Pgp becomes conformationally more dynamic, both in the TMDs and the NBDs, sampling two major conformations, termed TAR_*wide*_ and TAR_*closed*_, with different degrees of NBD dimerization (Fig. 2 and Fig. S1). Two major Pgp conformations, characterized by distinct degrees of NBD opening, were also observed from a recent inhibitor-bound cryoEM structures,^36,58^ with one of these structure showing an NBD distance within the range observed in our simulations for the TAR_*closed*_ conformations. One major finding of the present study is the lateral penetration of lipid molecules into the luminal region of Pgp, primarily from the cytoplasmic openings provided by the TMD portals. Lipid penetration into the lumen of Pgp takes place to different degrees in different bound states of Pgp, with the ATP systems showing the least and the TAR systems showing the largest number of such events. The differential lipid occupancy observed in the TAR_*closed*_ and TAR_*wide*_ (Fig. 5) may contribute to the small energy barrier between these two states that may be readily overcome through the diffusion of lipids from or into the binding cavity of Pgp.

Introduction of the inhibitor in the central cavity of Pgp, lying at the apex of the TMDs, leads to an increase in the fluctuations of neighboring protein residues (Fig. S4), as well as local unfolding of the portal helix TM10 (Fig. 3), one of the helices present at the drug entry Portal 2 and involved in forming direct interactions with the inhibitor. The zosuquidar-bound cryo-EM structure shows a similar disorder in TM10 in the presence of the inhibitor,^36^ although local unfolding in Portal 1 helices (TM4 and TM6) is also observed in this structure. This unfolding of the Portal 1 helices, which is not observed in other Pgp structures,^9,21,22^ has been suspected by a recent computational study^65^ to be an artifact of the small size of the nanodisc used in structural determination.

Compared to the transported substrates, Pgp inhibitors have been suggested to function by filling up the drug-binding cavity more completely, while at the same time, forming a larger number of contacts with the binding residues.^36,66^ Although the binding stoichiometry of TAR to Pgp is not well established, given its larger volume^9^ and flexibility (more than twice rotatable bonds as compared to zosuquidar for which an inhibitor-bound cryoEM structure exists^36^), only one TAR molecule may suitably fit inside the central cavity of Pgp. The single TAR molecule may instead successfully fill the binding cavity through recruiting lipids, as seen from the increased lipid density and lipid access events inside the central cavity of Pgp in our TAR-bound simulations compared to APO and ATP-bound systems (Fig. 4, Fig. 5 and Supplementary Video 1). Additionally, these “wedge-lipids”, where the lipid head groups from the cytoplasmic leaflet translate towards the membrane center while the lipid tails remain extended into the bulk membrane, are only observed in the inhibitor-bound systems (Fig. 6). The inhibitor may promote the recruitment of these wedge-lipids inside Pgp through a combination of conformational changes in the protein (larger TMD cytoplasmic portal/gate opening and local unfolding of TM10), encouraging lipid diffusion and translation, and then directly forming interactions with the lipid head groups inside the central cavity. It is possible that lipids binding to the central cavity of Pgp (in the absence of the inhibitor) may have similar effects on the dynamics of the transporter but to a lesser extent; in the absence of the inhibitors, lipid are presented less opportunities to penetrate the protein (e.g., due to more closed portals) and they may more readily diffuse out of the pocket.

A recent study using DEER spectroscopy suggested that high-affinity, third generation inhibitors like TAR can differentially modulate the interactions between the A-loop and ATP compared to substrate-bound Pgp, in turn inhibiting ATP hydrolysis. ^31^ A previous cysteine-scanning mutagenesis study has shown that the long-range conformational changes in the NBDs can be regulated by inhibitors/substrates.^62^ Additionally, lipid-binding sites observed at the regions formed by the kinking of the portal helices in response to inhibitor binding have been suggested to play a direct role in the modulation of Pgp function.^36^ Inhibitor-mediated lipid diffusion resulting in wedge-lipids inside the central cavity of Pgp, as observed in our simulations, may contribute to the differential coupling observed in the inhibitor-bound simulations compared to the APO and ATP systems (Fig. 7), modulating the transfer of signal between the TMDs and NBDs. This may in turn disrupt the natural dynamics of the NBDs, preventing their optimal interactions and dimerization and consequently inhibiting ATPase activity. In addition to directly restricting access to the substrate binding site, competitive inhibitors like TAR may thus successfully inhibit Pgp by stabilizing alternate occluded conformations, halting or shifting the equilibrium away from the OF states of the transporter.

The RMSF calculations evaluating differential fluctuations of Pgp in different bound states also highlighted that NBD2 is relatively more dynamic compared to NBD1 in the ATP-bound system (Fig. S4). This asymmetry in NBD dynamics, not observed in the APO system, is further amplified upon TAR binding. An asymmetric behavior in the NBDs of Pgp has also been reported in DEER studies of both ATP/substrate-bound and ATP/inhibitor-bound systems,^23,31^ where the asymmetry has been proposed to be a constitutive property of Pgp and directly involved in its asymmetric ATP hydrolysis and two-stroke transport mechanism of the OF state. Whether the conformational changes observed in our study in the IF state are correlated to the proposed transport mechanism in the OF state of Pgp remains to be further investigated. Additionally, as it is not possible to deconvolute the effect of ATP binding and TAR binding on the observed differential dynamics of the two NBDs in our simulations (both systems included ATP), we cannot determine whether these changes can be attributed to the chemical nature of the TAR molecule itself.

## Conclusion

As a multi-domain protein, Pgp displays a complex dynamical behavior, with the binding of ATP, substrates, inhibitors and surrounding lipids potentially modulating its conformational ensemble in many different ways. By employing extensive all-atom MD simulations, we show that a third generation, high-affinity inhibitor, TAR, allosterically modulates the conformational dynamics of both TMDs and NBDs, and their coupling, in Pgp, an additional mechanism that may contribute to its overall inhibitory effect on the transporter. (Fig. 8). This additional mode of inhibition, which is complementary to the inhibitor’s occupation of the substrate binding site (orthosteric inhibition), becomes amplified due to the inhibitor-mediated recruitment of wedge-lipids inside the central cavity of Pgp.

**Figure 8.**
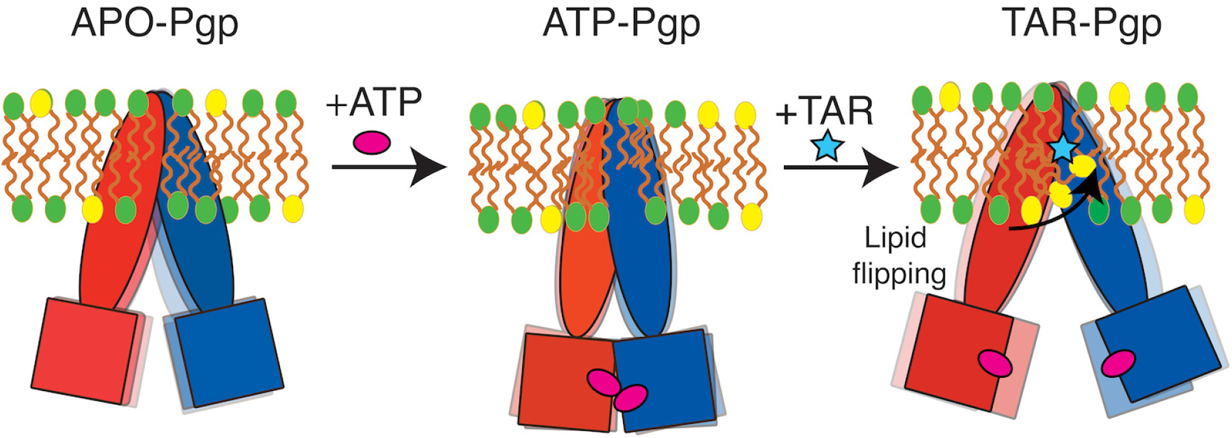
Modulation of the conformational dynamics of Pgp by the lipidic environment and bound inhibitor: In nucleotide-free (APO) state, Pgp exists in a dynamic equilibrium between open and closed IF conformations. ATP binding in the NBDs results in shifting of the equilibrium towards predominantly closed IF conformations of Pgp. The presence of a third-generation inhibitor (TAR) leads to increased lipid recruitment into the central cavity of Pgp, thereby, imposing a stronger inhibitory effect on NBD dimerization and closure of the cytoplasmic side of the TMDs.

Furthermore, we show that ATP binding to the NBDs (in the absence of inhibitor) modifies the conformational dynamics of the transporter, favoring closed IF conformations, which exist in a dynamic equilibrium with more open IF conformations. The open IF conformations captured for the APO simulations show large cytoplasmic opening that may allow active recruitment of substrate molecules, whereas the closed IF conformations favor the closure of cytoplasmic elements and, thereby, promote ATP binding and NBD dimerization.

To achieve the alternating access mechanism, membrane transporters are designed to undergo relatively larger conformational changes than other membrane proteins during their function. It is not therefore surprising that their structural dynamics and conformational changes can be significantly affected by membrane lipids. This mode of lipid effects might be more pronounced in ABC exporters such as Pgp in which large portals form by the extended helices in their structure, specially on the cytoplasmic side and in their IF state. In fact, alternate engagement and disengagement of lipids with the transporter should be considered almost critical parts/steps of the transport mechanism in these molecular machines. The coupling and interplay of these lipid events with ligand binding effects, reported in this study for the case of an inhibitor, can be expected to also exist for other modes of ligand binding and modulation in ABC transporters. One should expect ligand-mediated modulation of lipid-protein interactions even for the transported substrate, though in order for that transport mechanism to succeed the mode and extent of the interactions in this case has to be different from what we report here for a Pgp inhibitor.

From a pharmacological perspective, the additional role of lipids in inhibiting Pgp may also serve as a novel molecular strategy for the development of more potent and more specific inhibitors. The results presented in the study will provide another example of diverse mechanisms by which membrane lipids can modulate the function of complex membrane proteins such as Pgp.

## Acknowledgements

Research reported in this publication was supported by the National Institutes of Health under awards P41-GM104601 (to ET) and R01-GM123455 (to ET). We also acknowledge computing resources provided by Blue Waters at National Center for Supercomputing Applications, and Extreme Science and Engineering Discovery Environment (Grant MCA06N060 to ET). S.P. would like to acknowledge Beckman Institute Graduate Fellowship for funding.

