## Supplementary Figures for "Active Participation of Membrane Lipids in Inhibition of Multidrug Transporter P-Glycoprotein"

### Supporting Information

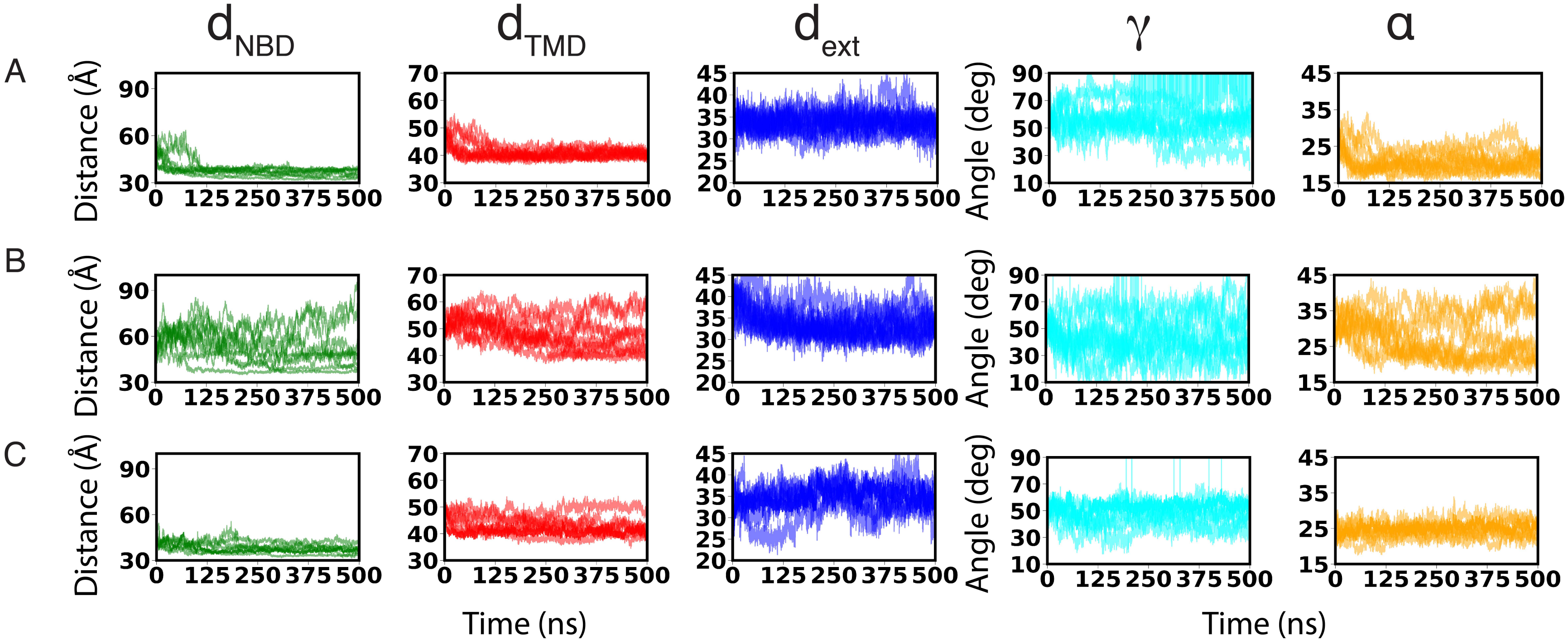


Figure S1: **Global conformational dynamics in Pgp:** Time evolution of the geometric parameters used to characterize the global conformational dynamics of Pgp for all the simulated replicas of (A) ATP, (B) TAR-bound, and (C) APO Pgp. The global conformational changes are monitored by $d_{NBD}$ (green), $d_{TMD}$ (red), $d_{ext}$ (blue), $\gamma$ (cyan), and $\alpha$ (orange).

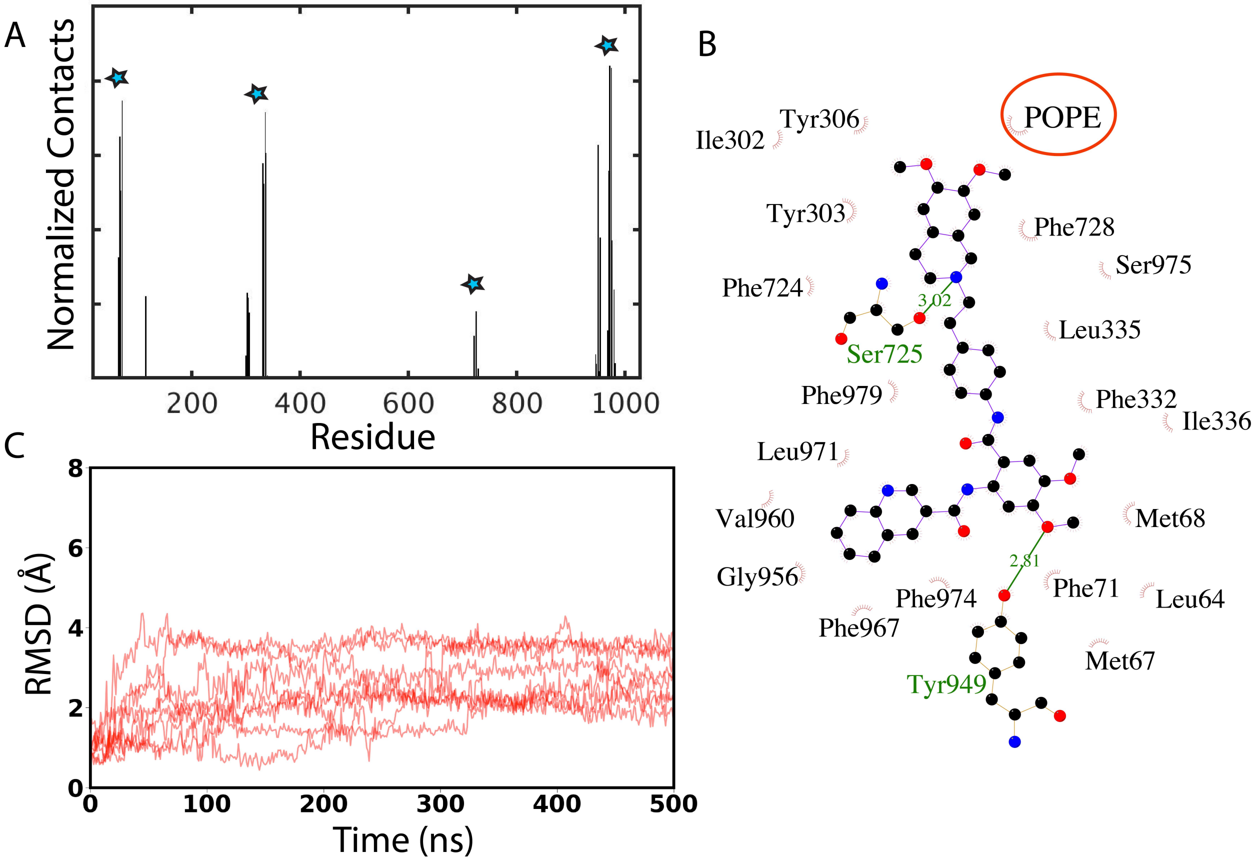


Figure S2: **TAR–Pgp interactions:** (A) All Pgp residues interacting with the inhibitor in a representative 500-ns simulation are shown. For comparison, the region of Pgp also found to be interacting with another third-generation inhibitor, zosuquidar, from a recent cryoEM structure (PDB:6FN1),[58] are shown with cyan stars. (B) Interaction of TAR with the surrounding residues of Pgp as captured in one of the simulations. The TAR binding pocket is predominantly rich in hydrophobic residues. Two polar side chains, Y949 and S725, form hydrogen bonds with the inhibitor. The position of a wedge-lipid POPE is shown in red circle. It should be noted that all the binding pocket residues present in the simulated mouse Pgp are conserved in human Pgp, except Ser725 which is mutated to Ala in human Pgp. (C) The stability of the bound TAR molecule was calculated by measuring the heavy atom RMSD of TAR in all 10 TAR-bound simulations. In all simulations, a stable mode of binding for the inhibitor was observed in the central cavity of Pgp.

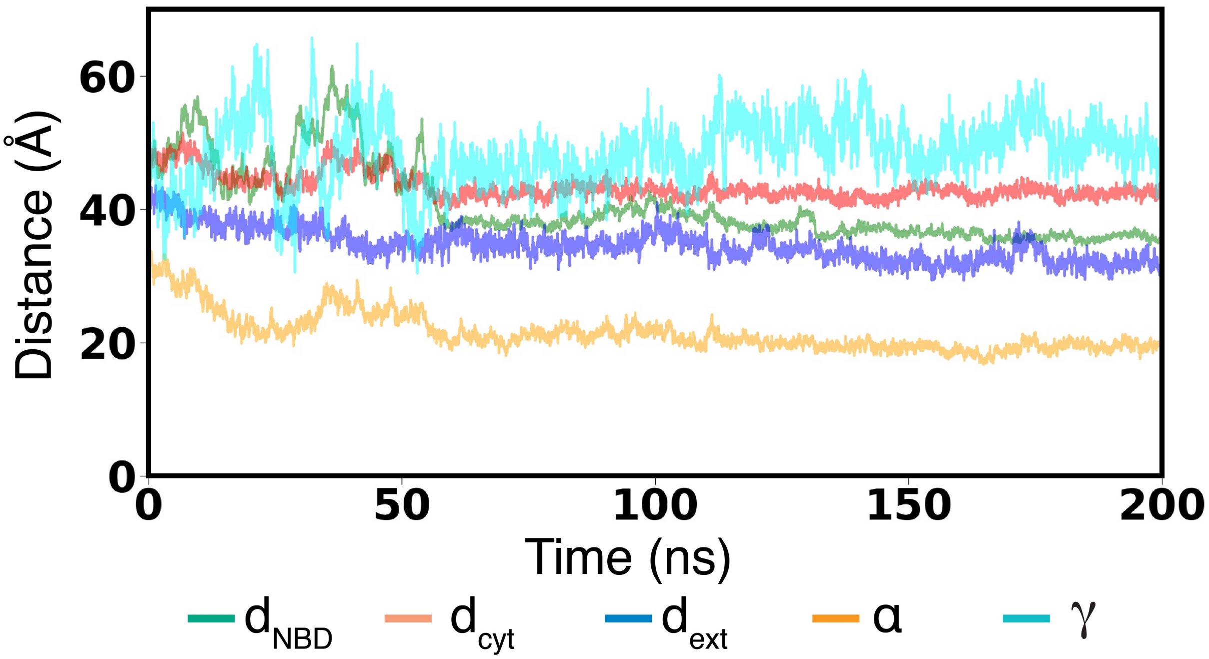


Figure S3: **Conformational dynamics of Pgp in the control (TAR-less) simulation:** Time evolution of the global conformation in TAR-bound Pgp conformation after removing the inhibitor is defined by $d_{NBD}$ (green), $d_{TMD}$ (red), $d_{ext}$ (blue), $\gamma$ (cyan), and $\alpha$ (orange). After removing TAR, the structure shows conformational dynamics similar to ATP-bound Pgp, with no lipid flipping events observed inside the central cavity.

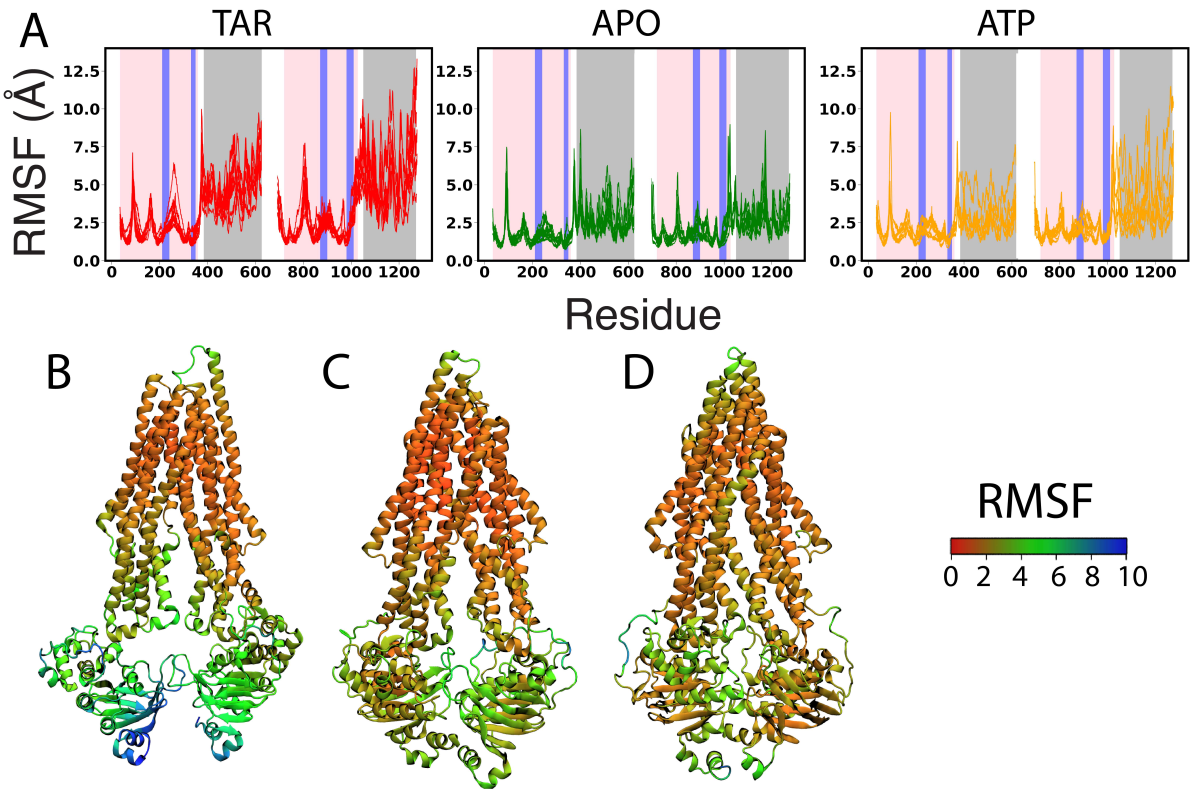


Figure S4: **Pgp fluctuation in MD simulations:** (A) RMSF calculated for all (10) the simulated systems. The RMSF values projected onto a representative molecular structure taken from respective simulations are shown for TAR-bound (B), ATP (C), and APO (D) Pgp. Scale bar for the RMSF values is shown on the right. Compared to ATP-bound and APO Pgp, overall higher RMSF values are observed in TAR-bound systems, especially in the portal helices and NBDs (highlighted with blue and grey backgrounds, respectively, in A). The TMD is highlighted in pink background.

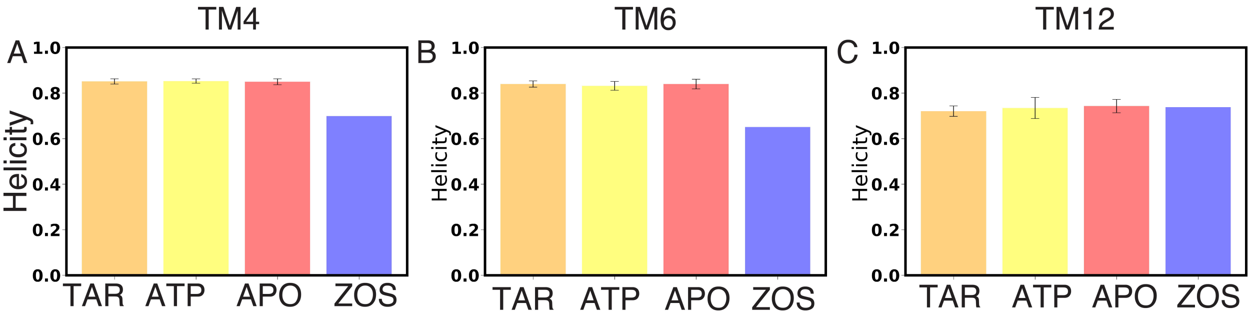


Figure S5: **Local changes in secondary structure:** Comparison of the degrees of helicity for portal helices TM4 (A), TM6 (B), and TM12 (C) in APO, ATP, and TAR-bound Pgp with the zosuquidar-bound cryoEM structure (ZOS, blue).[58] For all three portal helices, similar degrees of helicity was observed in all three simulated systems. As compared to zosuquidar-bound Pgp, no loss in helicity was observed for TM4 and TM6.

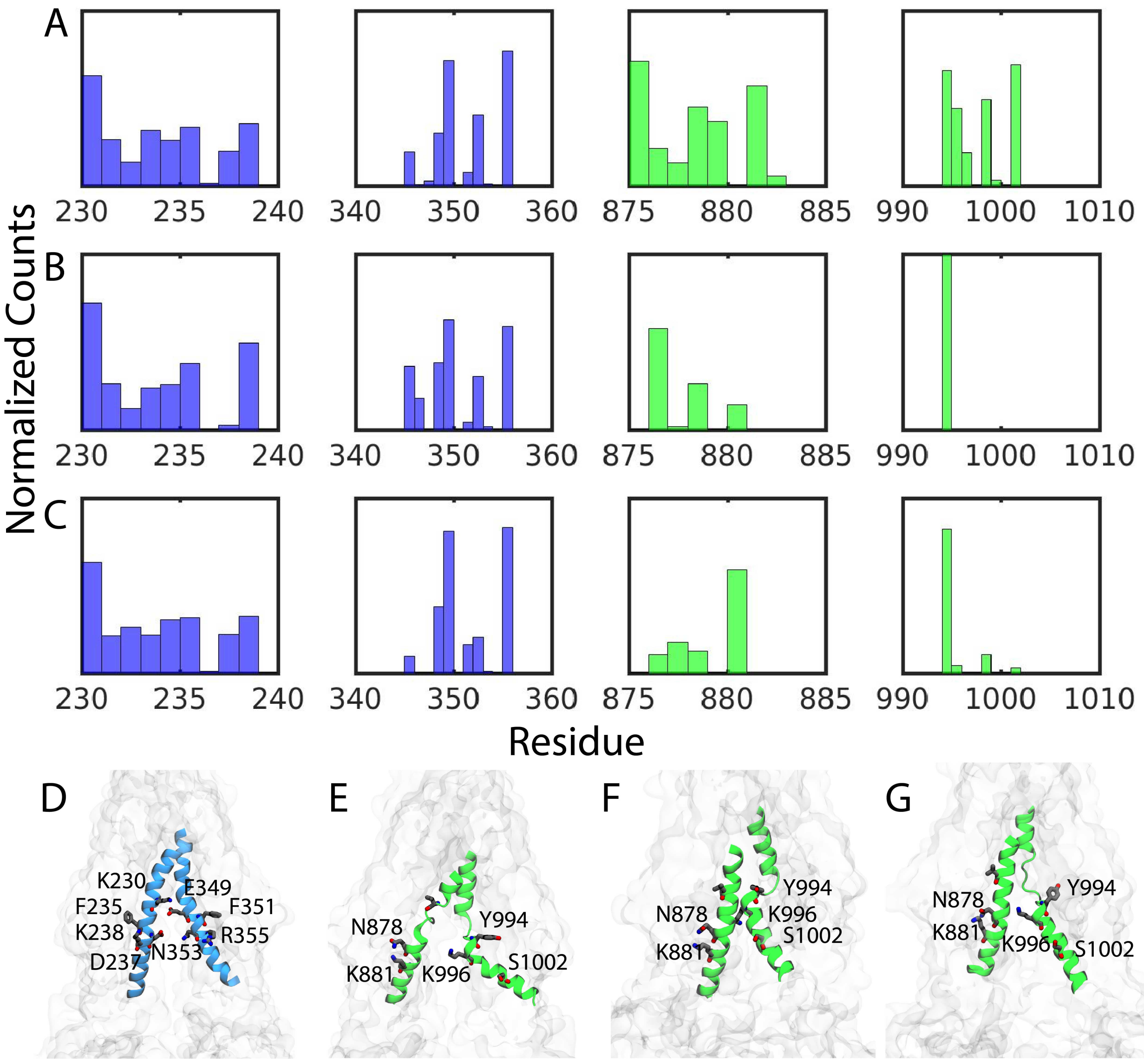


Figure S6: **Lipid-interacting residues at the entry portals:** Residues in Portals 1 and 2, shown in blue and green, respectively, making contacts with penetrating lipids over the course of the simulation in TAR-bound (A), ATP (B), and APO Pgp (C). As compared to Portal 1, less lipid contacts are observed in Portal 2. All the portal residues making contact for more than 20% of the simulation time are shown for Portal 1 (not undergoing structural change) (D) and for Portal 2 (undergoing differential unfolding in TM10 helix) in (E) TAR-bound, (F) ATP-bound, and (G) APO systems. In addition to the differences in the overall opening of Portal 2, we also observed differential side-chain arrangements at this portal, in different simulated systems.

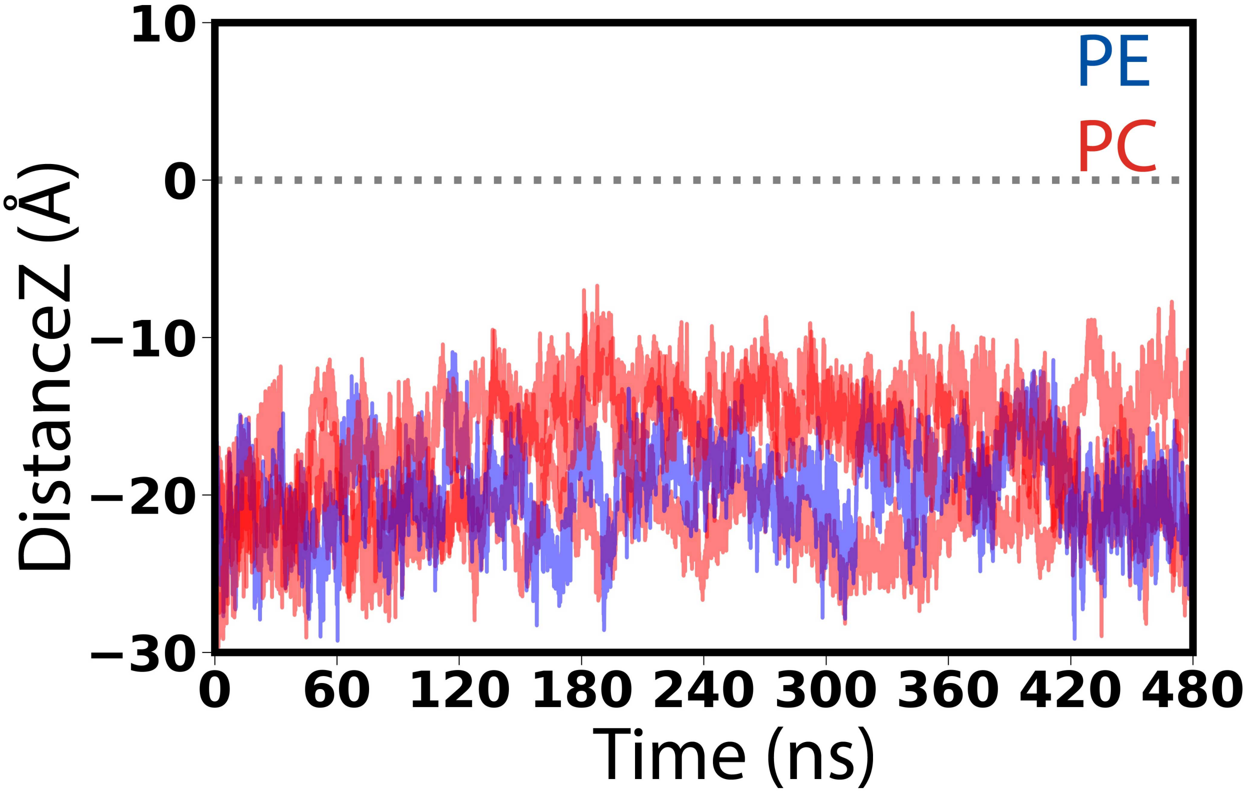


Figure S7: **Monitoring lipid head-groups inside the lumen of APO Pgp:** Lipid head-group movement in the central cavity of Pgp was monitored by calculating the $z$ component of the center of mass of the lipid head-group in all the simulation replicas. The midplane ($z = 0$) of the lipid bilayer is marked by a dashed line. Overall, we did not observe any head-groups flipping (crossing $z = 0$, as observed, e.g., in the TAR systems).

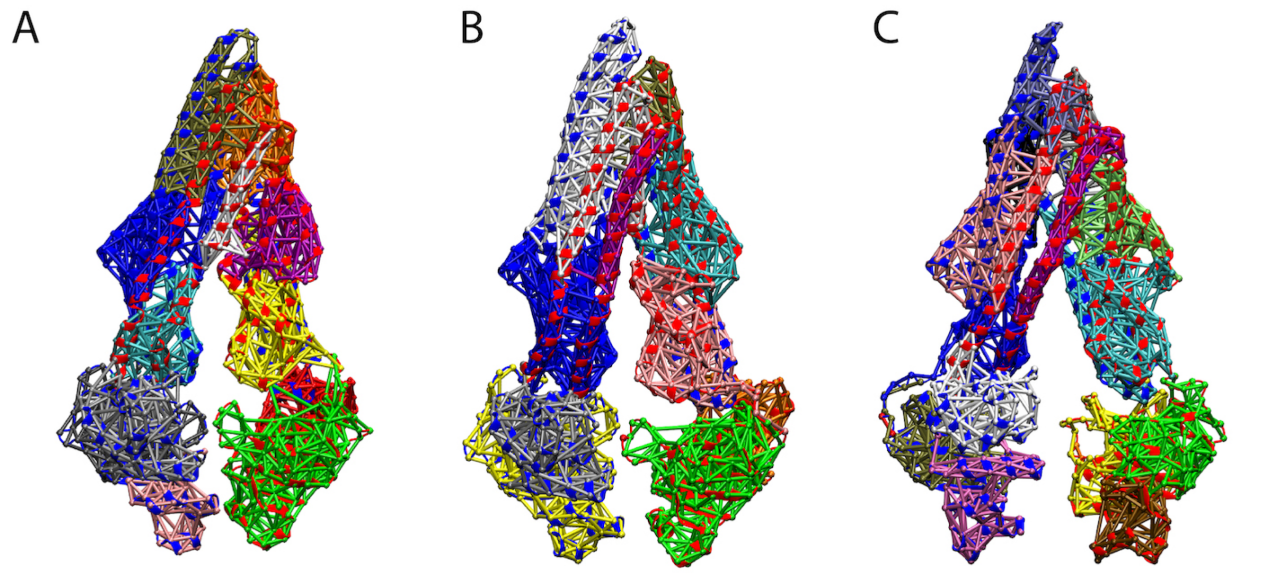


Figure S8: **Network-based community analysis:** Community analysis in the APO (A), ATP-bound (B), and TAR-bound (C) Pgp showing regions with high correlation. Analysis was performed on a representative, 500-ns MD simulation replica for each system. The communities are individually colored and show similar behaviors in all three systems.
